# A novel splice variant of human TGF-β type II receptor encodes a soluble protein and its Fc-tagged version prevents liver fibrosis *in vivo*

**DOI:** 10.1101/2021.03.01.433173

**Authors:** Marcela Soledad Bertolio, Anabela La Colla, Alejandra Carrea, Ana Romo, Gabriela Canziani, Stella Maris Echarte, Sabrina Campisano, German Patricio Barletta, Alexander Miguel Monzon, Tania Melina Rodríguez, Andrea Nancy Chisari, Ricardo Alfredo Dewey

**Affiliations:** Laboratorio de Terapia Génica y Células Madre, Instituto Tecnológico de Chascomús (INTECH), CONICET-UNSAM), Chascomús, Buenos Aires, Argentina (B7130); Departamento de Química y Bioquímica, Facultad de Ciencias Exactas y Naturales, Universidad Nacional de Mar del Plata, Buenos Aires, Argentina (B7602AYL); Drexel U-Sidney Kimmel Cancer Center / Thomas Jefferson U S200 Biosensor Shared Resource, Department of Biochemistry and Molecular Biology, Drexel University College of Medicine, Philadelphia, USA (PA19102); Molecular Physics and Biophysics Group, Department of Science and Technology, National University of Quilmes, CONICET, Argentina; Department of Biomedical Sciences, University of Padua, Via Ugo Bassi 58/B, Padua, 35121, Italy

**Author notes:** Corresponding author: Prof. Ricardo Alfredo Dewey, Laboratorio de Terapia Génica y Células Madre, Instituto Tecnológico de Chascomús (INTECH), CONICET-UNSAM, (B7130) Chascomús, Buenos Aires, Argentina. Centro Regional de Estudios Genómicos (CREG), Universidad Nacional de La Plata, La Plata, Argentina. These authors contributed equally (Marcela Soledad Bertolio, Anabela La Colla, Alejandra Carrea).

## Abstract

We describe, for the first time, a new splice variant of the human TGF-β type II receptor (TβRII). The new transcript lacks 149 nucleotides, causing a frameshift with the appearance of an early stop codon, rendering a truncated mature protein of 57 amino acids. The predicted protein, lacking the transmembrane domain and with a distinctive 13 amino acid stretch in the C-terminus, was named TβRII-Soluble Endogenous (TβRII-SE). Binding predictions indicated that the novel 13 amino acid stretch interacts with all three TGF-β cognate ligands and generate a more extensive protein-protein interface than TβRII. TβRII-SE and human IgG1 Fc-domain, were fused in frame in a lentiviral vector (Lv) for further characterization. With this vector, we transduced 293T cells and purified TβRII-SE/Fc by A/G protein chromatography from conditioned medium. Immunoblotting revealed homogeneous bands of approximately 37 kDa (reduced) and 75 kD (non-reduced), indicating that TβRII-SE/Fc is secreted as a disulphide-linked homodimer. Moreover, high affinity binding of TβRII-SE to the three TGF-β isoforms was confirmed by Surface Plasmon Resonance (SPR) analysis. Also, intrahepatic delivery of Lv.TβRII-SE/Fc in a carbon tetrachloride-induced liver fibrosis model revealed amelioration of liver injury and fibrosis. Our results indicate that TβRII-SE is a novel member of the TGF-β signaling pathway with distinctive characteristics. This novel protein offers an alternative for the prevention and treatment of pathologies caused by the overproduction of TGF-β ligands.

## Introduction

Transforming growth factor-β (TGF-β) is a multifunctional cytokine involved in critical processes, such as cell proliferation, maturation, and differentiation, wound healing, and immune regulation.^1^ Three TGF-β isoforms have been identified in mammals: TGF-β1, TGF-β2 and TGF-β3. They share 64–82% sequence identity and are encoded by distinct genes that are expressed in developmentally regulated and tissue-specific manners.^2^ Although the phenotypes of isoform-specific null mice are non-overlapping,^3-5^ indicative of distinct temporal-spatial roles *in vivo*, the ligand isoforms exhibit overlapping biological activities in tissue culture assays.

Canonical signaling starts when mature, dimeric TGF-β family ligands bind cell surface receptor complexes that combine structurally related transmembrane kinases: two !type II” (TβRII) and two !type I” (TβRI) receptors.^6^ TβRI has a short Gly-Ser–rich juxtamembrane sequence (GS domain), that is phosphorylated by TβRII kinase in response to ligand binding.^6-8^ Not all TGF-β ligands contact the receptor complex equally. TGF-β1 and TGF-β3 bind with high affinity to TβRII receptor dimers without the need for TβRI receptors, whereas TβRI receptors have a low affinity for TGF-β and require TβRII for ligand binding.^9^ Conversely, TGF-β2 has low affinity for both the TβRII or TβRI receptors suggesting a process with TGF-β2 initially binding to either the TβRI or TβRII receptors in preformed receptor complexes.^10^ Moreover, it is known that the betaglycan coreceptor (former TβRIII), is necessary for efficient binding of TGF-β2 and subsequent signaling.^11^ This co-receptor binds all three TGF-βs with a highest affinity for TGF-β2.^12^ In this model, TGF-β2 binds to betaglycan, which then recruits TβRII and TβRI resulting in phosphorylation of TβRI and downstream signaling.^11^

Ligand binding to the receptor ectodomains induces conformational changes at the ligand-receptor interface, bringing their cytoplasmic domains closer^7,9^ This stabilization enables TβRII to phosphorylate TβRI at the serine residues of the GS domain, which then induce conformational changes, that activate TβRI kinase.^7,13^ TβRI activated by TβRII, in turn, activate effector Smads (Smad2 and 3) through phosphorylation of their two C-terminal serines. These !receptor-activated Smads” (R-Smads) then combine with Smad4 to form complexes that translocate into the nucleus, where they activate or repress target genes in cooperation with high-affinity DNA binding transcription factors and coregulators.^14^

In addition to TβRII protein, the *tgfbr2* gene encodes the membrane-anchor isoform TβRII-B, via alternative splicing. This variant involves an insertion of 75 bp coding for 25 amino acids in the extracellular domain of the receptor, with an isoleucine to valine exchange.^15^ Unlike TβRII, TβRII-B binds and signals directly via all three TGF-β isoforms without the requirement of betaglycan.^16^ Additionally, in the absence of betaglycan, TGF-β2 binding to TβRII-B requires TβRI.^17^

The canonical signaling model described above is an oversimplification that does not explain the highly diverse and context-dependent TGF-β responses.^1,18^ Also, TGF-β receptors activate non-Smad signaling pathways that substantially contribute to the TGF-β response, including the MAPK and phosphoinositide 3-kinase (PI3K)–AKT–mammalian target of rapamycin (mTOR) pathways,^19^ adding more complexity to the signaling cascade. It has become apparent that much remains to be learned about the mechanisms involved in receptor presentation and activation, and in the control of cell responsiveness to define the developmental and pathological roles of TGF-β signaling.^18^

Adding another turn to the TGF-β signaling pathway, here we describe, for the first time, TβRII-SE, a novel human TβRII splice variant which encodes a truncated soluble isoform of the receptor. Opposite to TβRII and its splice variant TβRII-B, Tβ-RII-SE isoform binds all three TGF-β ligands, in the picomolar affinity range, without participation of additional receptors.

Given to the enhanced TGF-β1 signaling in cancer and fibrosis,^1^ TGF-β has become a promising therapeutic target. Here, we checked TβRII-SE functionality in a liver fibrosis animal model. Liver fibrosis is a common stage of all chronic liver diseases (CLD)^20^ and is characterized by the excessive synthesis and accumulation of extracellular matrix proteins (ECM). After liver damage, reparative mechanisms are activated to replace injured hepatocytes. However, if the exposure to the liver injury agent persists over a long time, the continuous wound healing response leads to the destruction of liver architecture and, eventually, results in liver cirrhosis, liver failure and high risk to develop hepatocarcinoma (HCC) .^21^ Even though for many years it was thought that liver fibrosis was irreversible and relentlessly progressive, strong evidence has shown that, on the contrary, that liver fibrosis is a highly dynamic and reversible process.^22^ TGF-β plays an important role during all phases of the development of liver fibrosis, being responsible for hepatic stellate cells (HSC) activation to myofibroblasts (MFB),^23^ reactive oxygen species (ROS) generation,^24^ and ECM production stimulation.^25^ Therefore, to functionally evaluate the capacity of TβRII-SE to modulate TGF-β effect *in vivo*, we constructed a lentiviral vector encoding TβRII-SE fused in frame with IgG1 Fc domain. Intrahepatic infusion of this vector, in a carbon tetrachloride (CCl_4_)-induced liver fibrosis model, suggested a strong protective effect of TβRII-SE against liver fibrogenesis.

## Results

### TβRII-SE is a novel TβRII-splice variant

The new TβRII-splice variant was first identified as a 433 bp fragment by end point RT-PCR in human peripheral blood-derived T lymphocytes (Fig. 1a). The primer pair employed also amplified the cDNA sequence encoding the ER-signal peptide (SP), the extracellular domain (ECD), and the transmembrane domain (TMD) corresponding to the two membrane bound TβRII-splice variants (TβRII and TβRII-B) (Fig. 1b). Using this primer pair, we also found the presence of the 433 bp band in CD3^+^, CD19^+^, CD14^+^ cells isolated by immunomagnetic separation, and in granulocytes obtained from Ficoll density gradient (Fig. 1c). In addition, the new band was also also detected in primary cultures of adipose derived mesenchymal stromal cells (hASC), and in the cell lines 293T and Jurkat (Fig. 1c).

**Figure 1.**
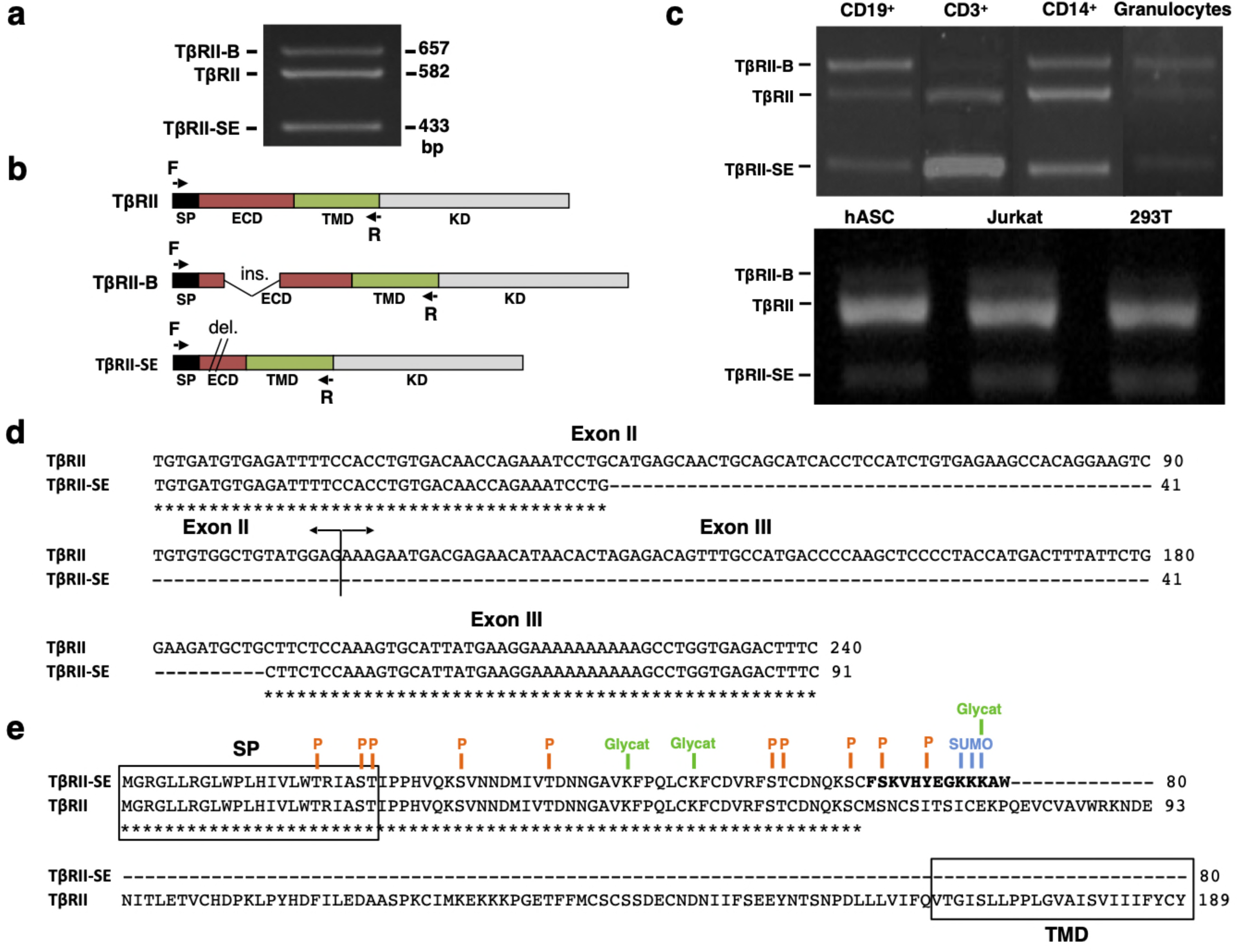
TβRII-SE splice variant is produced by human cells and is able to generate a truncated soluble protein. **a** TβRII-SE was originally detected in human lymphocytes as a 433 bp band after end point RT-PCR. **b** Primers design to amplify the coding sequence of the endoplasmic reticulum signal peptide (SP), extracellular domain (ECD) and transmembrane domain (TMD) of membrane bound TβRII splice variants. **c** End point RT-PCR of PBMC isolated CD3^+^, CD19^+^, CD14^+^ cells, and granulocytes; and human adipose stromal cells (hASC), immortalized T lymphocyte cell line (Jurkat), and human embryonic kidney cell line containing the SV40 T-antigen (293T). **d** Partial nucleotide sequence alignment of TβRII and Tβ-RII-SE depicting the stretch absent in the new splice variant. **e** Predicted amino acid sequence alignment of TβRII and TβRII-SE showing the presence of the SP and predicted PTM in both isoforms, and the absence of the transmembrane domain (TMD) in TβRII-SE. C-terminal 13 amino acid stretch distinctive of TβRII-SE is shown in bold. P= kinase-specific phosphorylation; Glycat=glycation; and SUMO= non-consensus sumoylation.

In comparison to TβRII, DNA sequence analysis revealed, in the new PCR amplified fragment, the absence of 149 bp (63 bp of exon II and 86 bp of exon III), indicating alternative splicing (Fig. 1d). The predicted amino acid (AA) sequence of this new splice variant showed that the lack of 149 bp alters the coding sequence, creating a small open reading frame (sORF) with a premature stop codon (Fig. 1e). Therefore, the new TβRII splice variant has the capacity to encode a protein of 80 AA residues, which includes an ER signal sequence of 23 AA, and a mature protein of 57 AA, lacking the transmembrane domain (TMD) (Fig. 1e). Thus, we named the novel TβRII splice variant as TβRII soluble endogenous or TβRII-SE. TβRII and TβRII-SE protein sequence alignment revealed, in the C-terminus of the new isoform, a stretch of 13 AA (FSKVHYEGKKKAW) distinctive of TβRII-SE (Fig. 1e). Post-translational modification (PTM) predictions of this novel receptor indicated a putative glycation site (K55), kinase specific phosphorylation (S46 and Y50), and non-consensus sumoylation sites (K53, 54 and 55) (Fig. 1e and Table S1). The predicted molecular weight of the mature TβRII-SE, without PTM, was 6532.51 Da with isoelectric point (pI) of 9.05 (Table S1).

### TβRII-SE splice variant encodes an stable peptide with distinctive binding properties

Based on TβRII-SE predicted AA sequence, we generated a 3D model using the Robetta server, and the TβRII extracellular (EC) domain as a template (Fig. 2a). The stability of the model was confirmed with a long molecular dynamics trajectory. This 3D modeling revealed that TβRII-SE cysteines would not be able to form disulphide bonds as in TβRII-EC (Fig. 2a). In addition, to predict how TβRII-SE binds to TGF-β ligands, we superimposed the mature protein model of the new receptor isoform over the crystallographic structure 2PJY, corresponding to the TβRII/TβRI/ TGF-β3 complex. In this analysis we found that TβRII-SE and TGF-β3 establish stable physical contacts, in particular, lysine 53 (K53) of TβRII-SE forms cation-Pi interaction with tryptophan 32 (W32) of TGF-β3, and phenylalanine 60 (F60) of TβRI, whereas tryptophan 57 (W57) of TβRII-SE makes an intramolecular stacking with phenylalanine 24 (F24), promoting a β-fork structure (Fig. 2b Upper right panel and Fig. 2c). Similarly, histidine 49 (H49) and tyrosine 50 (Y50) of TβRII-SE form an aromatic stacking with histidine 34 (H34) and tyrosine 91 (Y91) of TGF-β3 (Fig. 2b lower right panel and Fig. 2c). Notably, glutamic acid 51 (E51) of TβRII-SE can form salt bridges with arginine 25 (R25) of TGF-β3, substituting the same interaction between glutamic acid 119 (E119) in the TβRII-EC (extracellular) domain. The interactions present in the distinctive 13 AA stretch of TβRII-SE allow to share with TβRII the same TGF-β binding region (Fig. 2c), although both TβRII isoforms are structurally different. These new interactions suggested a more extensive protein-protein interface than TβRII, allowing TβRII-SE to compete for the binding with TβRI and TGF-β ligands, in the trimeric complex.

**Figure 2.**
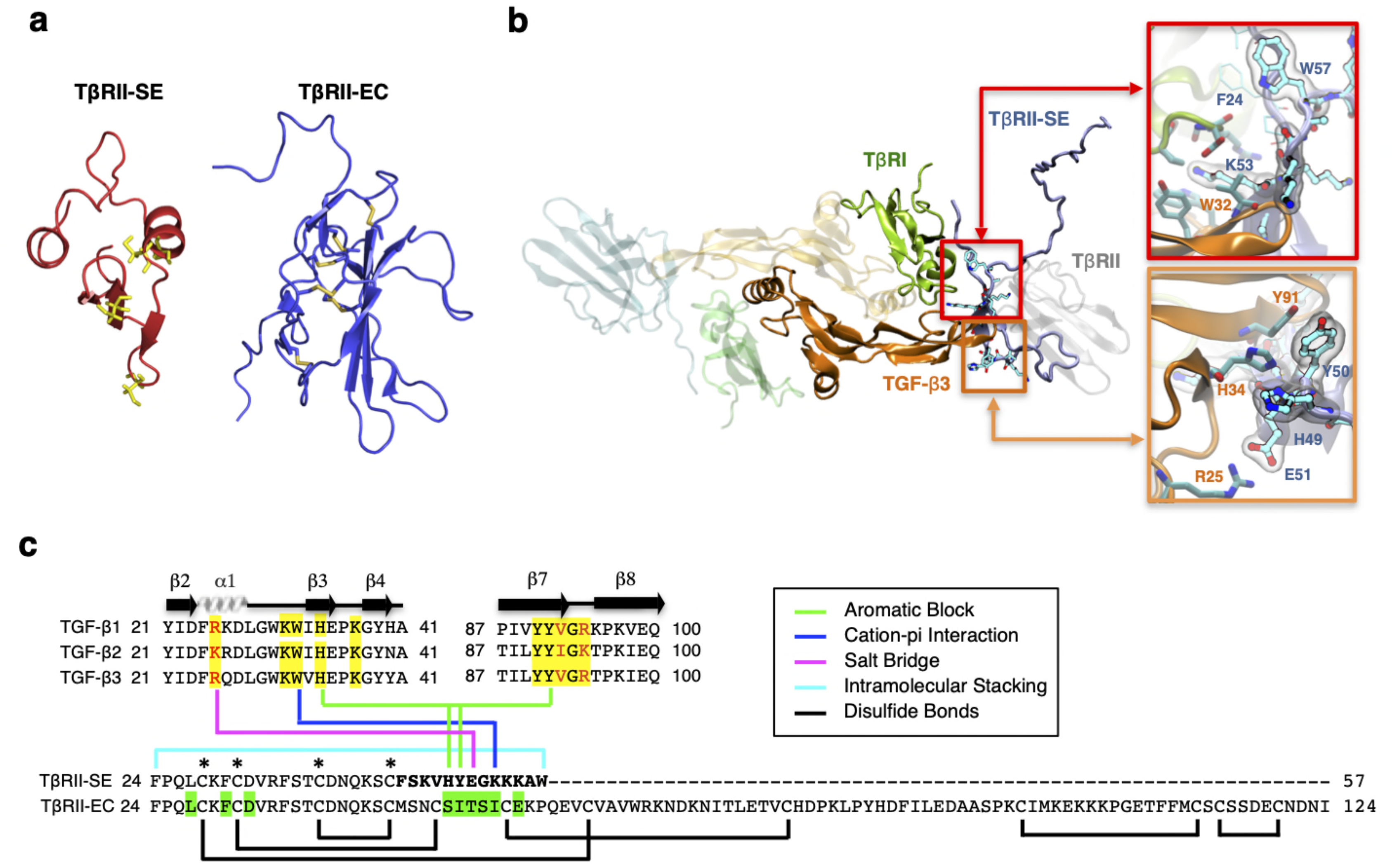
3D modeling reveals unique characteristics of TβRII-SE in comparison to the extracellular domain of membrane bound TβRII. **a** TβRII-SE and TβRII-EC 3D models showing cysteine residues in yellow, and lack of disulphide bonds in Tβ-RII-SE. **b** 3D model of mature TβRII-SE superimposed over the crystallographic structure 2PJY (TβRII/TβRI/TGF-β3) (left panel). Amino acids interaction between TβRII-SE (blue residues) and TGF-β3 (orange residues) are shown magnified in the right upper and lower panels. **c** Amino acid sequence alignment of human TGF-β family members (top panel) showing residues interacting with TβRII-EC (yellow shading), and residues interacting with TβRII-SE (colour lines linking the bottom panel). Green shading indicates residues in TβRII-EC interacting with TGF-β cognate ligands. Brackets and asterisks indicate cysteine residues forming and not forming disulphide bonds in TβRII-EC and TβRII-SE, respectively.

### TβRII-SE Fc-tagged over expression *in vitro*

To generate high levels of TβRII-SE protein for further studies, we aimed to enhance transgene expression in mammalian cells by modifying TβRII-SE coding sequence by inclusion of a Kozak sequence and codon optimization (co) (Fig. S1). In addition, to ease protein purification and to increase serum half life of the peptide, we fused coTβRII-SE to human IgG1-Fc domain coding sequences “in frame”, and cloned it into a lentiviral vector to make Lv.TβRII-SE/Fc (Fig. 3a).

**Figure 3.**
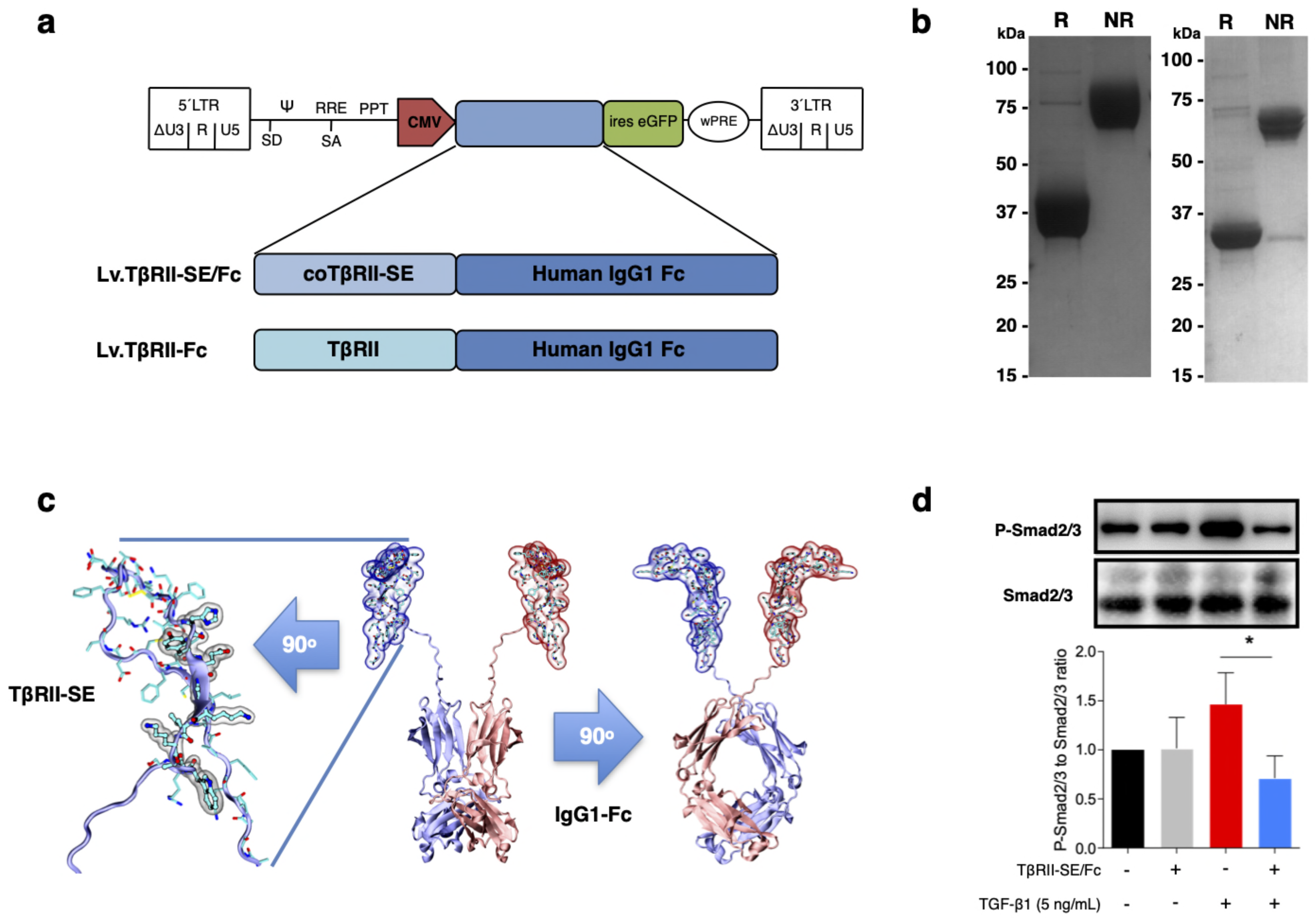
TβRII-SE/Fc overexpressed in human cells is secreted to the extracellular milieu as a disulphide-linked homodimer, and inhibits Smad2/3 activation. **a** Either coTβRII-SE/Fc or TβRII-EC were ligated to the human IgG1 Fc coding sequence and cloned into a self inactivating (SIN) bicistronic lentiviral vectors making Lv.TβRII-SE/Fc and Lv.TβRII-Fc, respectively. The proviral lentiviral vectors contain long terminal repeat sequences (LTRs) on both ends, devoid of LTR promoter/enhancer sequences (ΔU3) after reverse transcription. The cytomegalovirus (CMV) promoter was used as internal promoter to drive the expression of either TβRII-SE/Fc or TβRII-Fc. Furthermore, the vectors include the safety-improved woodchuck hepatitis virus post-transcriptional regulatory element (wPRE), a splice donor (SD), a splice acceptor (SA) and the packaging signal (Ψ), together with the Rev responsive element (RRE), and the central polypurine tract (PPT). **b** SDS-PAGE and Coomassie blue staining of Protein A/G purified supernatants of TβRII-SE/Fc overexpressing 293T cells under reducing (R) and non-reducing (NR) conditions. **c** Dimeric 3D model of TβRII-SE/Fc using the structures 2PJY (TβRII) and 1L6X (Fc), as templates (middle panel). TβRII-SE is shown magnified and rotated 90° (left panel) with atomic representation wrapped in its solvent-accessible surface area. Amino acids with solvent-accessible surface (highlighted in grey) are part of the protein-to-protein interface. Also, TβRII-SE/Fc 3D model is shown rotated 90° (right panel). **d** Representative immunoblotting (top panel) and semi-quantitative densitometry of three independent experiments (bottom panel) of P-Smad2/3 and total Smad2/3 in HCT116 cells overexpressing TβRII-SE/Fc (+) and control (-), in the presence (+) and absence (-) of TGF-β1 stimulation. *p<0.05.

We next overexpressed TβRII-SE/Fc chimera by means of a lentiviral vector (Lv.-TβRII-SE/Fc), in 293T cells. Recombinant protein purified by A/G protein chromatography from 293T cells conditioned medium was analyzed by SDS-PAGE gels. Coomassie Blue staining revealed, under reducing and non-reducing conditions, broad bands of approximately 37-38 kDa and 75 kD, respectively (Fig. 3b left). We also overexpressed TβRII-SE/Fc without N-glycosylation in the CH2 domain of the Fc (Fig. 3b right). Here we observed bands of 32 kDa and 64 kDa under reducing and non-reducing conditions, respectively. These data were consistent with TβRII-SE/Fc being secreted as a disulphide-linked homodimer, as predicted by a 3D dimeric model (Fig. 3c). To determine the potential role of TβRII-SE on downstream TGF-β signaling, we tested its effect on Smad2/3 activation. Lysates from HCT116 cells either transduced with Lv.TβRII-SE/Fc or untransduced (control), in the presence or absence of TGF-β1, were subjected to both, anti-Smad2/3 and anti-pSmad2/3 immunoblotting (Fig. 3d). TGF-β1 induced Smad2/3 activation in control HCT116 cells, but not in TβRII-SE/Fc overexpressing cells. These results suggested that TβRII-SE is able to block TGF-β signaling cascade by trapping TGF-β1, as suggested by 3D modeling.

### TβRII-SE/Fc binds with high affinity to all three TGF-β isofoms

We next examined TGF-β1, 2 and 3 binding to TβRII-SE/Fc using surface plasmon resonance (SPR). This is a label-free method that was optimized to rank the kinetics of interaction of the three TGF-β isoforms with TβRII-SE relative to the native ligand, the TβRII receptor, and the pan-isoform specific 1D11 MAb, a TGF-β neutralizing antibody with therapeutic efficacy and characterized kinetics of binding. MAb 1D11 binds the three TGF-β isoforms with fast association and slow dissociation kinetics, forming stable homodimers. To assess the kinetics of potentially tight TGF-β-TβRII-SE/Fc complexes, interactions were measured at flow 50 µL min^-1^ using 60 to 90-seconds associations, 15-minute dissociations and low ligand surface densities. We observed rapid and stable kinetics of binding of TGF-β isoforms with TβRII-SE, TβRII and 1D11 MAb with the exception of TGF-β2 with TβRII, previously described to form fast dissociating complexes. We note that 1:1 interactions that dissociate at rates slower than 1×10^−3^ s^-1^ require either longer association phases or higher concentrations to reach steady state. Consistent with the short associations, and to resolve kinetic differences among TGF-β, we monitored binding at concentrations that were 250-fold the previously reported K_D_s of complexes with TβRII-Fc and 1D11, to allow at least one of the TGFβ isoforms binding to reach steady state in 90 seconds. TβRII-Fc binding TGF-β isoforms reached steady state within 90 seconds, particularly the lower affinity TGF-β2 which was not measured to ligand saturation, while TβRII-SE/Fc and 1D11 complexes reached steady state at saturation for at least one isoform, all interactions displaying slow complex dissociation phases (*k*_*d*_*s* ranging 2-10 ×10^−4^ s^-1^). Association and dissociation rate constants (*k*_*a*_ and *k*_*d*_) for each TGF-β1, TGF-β2 and TGF-β3 were best estimated using a single step, 1:1 binding model for the two-site binding reactions which were predicted to be simultaneous with TβRII-SE and Tβ-RII receptor pairs brought into close proximity by the Fc-sequence dimerization. Binding profiles for 30 nM TGF-β 1, 2 and 3 to TβRII-SE-Fc are overlayed and compared with those obtained with TβRII-Fc and 1D11 in Fig. 4a, b and c, respectively. Fast *k*_*a*_s ranging 1 to 10 ×10^6^ M^-1^s^-1^, slow to medium *k*_*d*_s ranging 2 to 10 ×10^−4^ s^-1^, and maximum binding capacities or R_max_ were calculated for all replicate interactions (1:1 model fit, Chi2 <0.02*Rmax).

**Figure 4.**
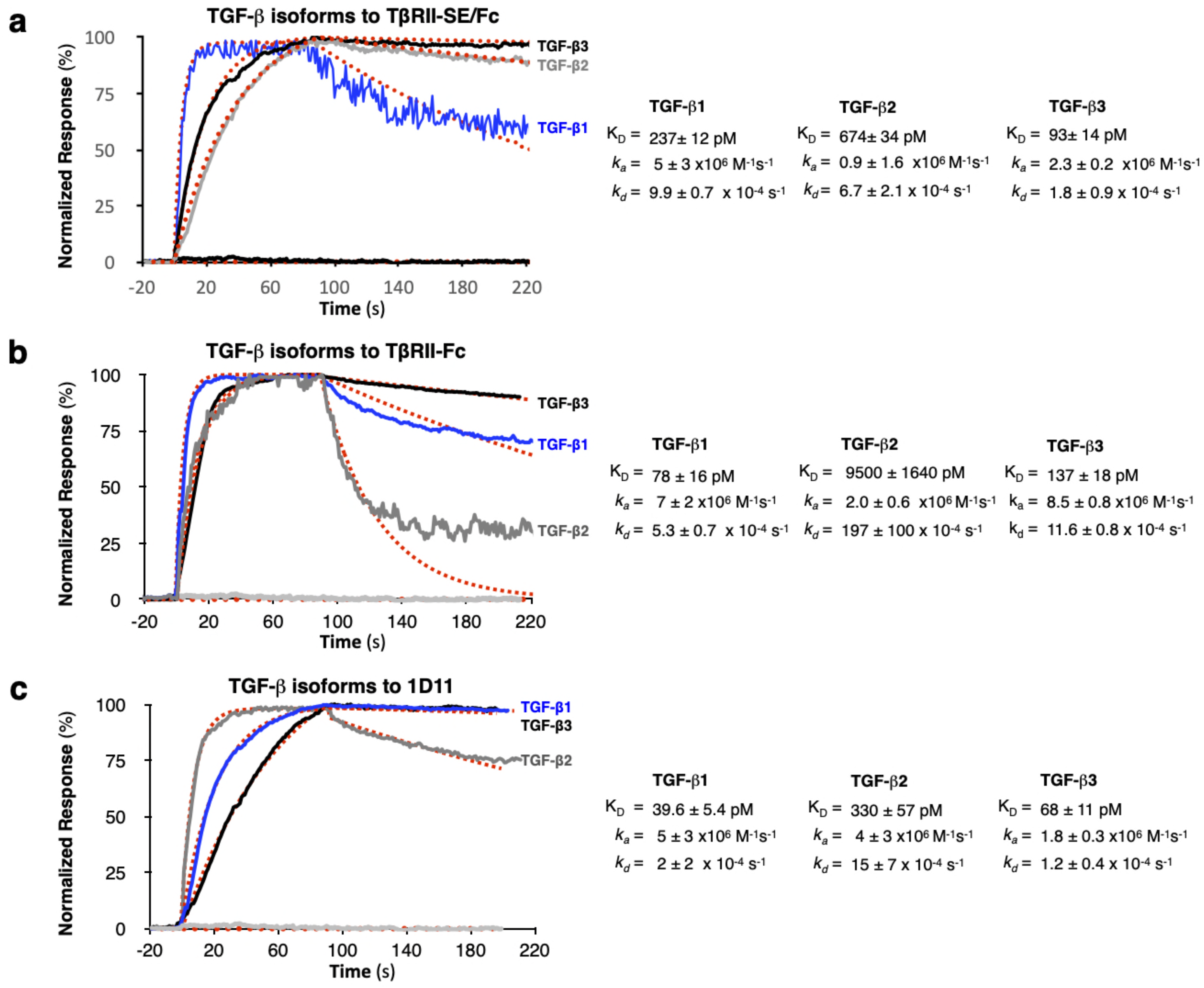
SPR binding profiles of 30 nM TGF-β isoforms to Fc-captured recombinant TβRII-SE/Fc, TβRII-Fc, and 1D11 mAb. Normalized association (t =90s) and dissociation profiles (t =130s shown) for **a** TβRII-SE/Fc, **b** TβRII-Fc, and **c** 1D11 mAb complexes with each TGF-β isoform are shown overlayed with the respective global model fits (red dashed lines). Blue: TGF-β1, grey: TGF-β2 and Black: TGF-β3. Calculated kinetic parameters are shown to the right for each interaction.

Stoichiometries of complex formation were calculated to explore the mechanistic dominance of homodimeric TGF-β isoforms binding with TβRII-SE/Fc. Surface densities (RU/KDa molecular mass) ranged from 0.6-2.4 and resulted in maximum signals (R_max_) ranging 45-126 RU for each ligand due to equal accessibility to all binding sites. Maximum binding capacities (or Rmax) were used to calculate the stoichiometries of binding after each ligand capture was optimized and ligands were not subjected to any regeneration steps that reduce binding capacity. Supplemental Table 2 shows the theoretical versus the experimental stoichiometries of binding and was estimated as one TGF-β dimer per TβRII and TβRII-SE bivalent Fc ligand, and two TGF-β dimers per 1D11 as was previously reported in Fab-fragment binding of other neutralizing antibodies that functionally mimic the binding mode of both TβRII and TβRI receptors to the dimer interface. Taken together, these results indicate that TβRII-SE/Fc binds all three TGF-β isofoms with sub-nanomolar affinities that were comparable to the affinities of their complexes with 1D11, but with stoichiometry similar to those of TβRII-Fc binding.

### TβRII-SE/Fc overexpression prevents CCl_4_-induced liver fibrosis

We next aimed to evaluate the functionality of TβRII-SE/Fc *in vivo*, in comparison to TβRII-Fc, in a CCl_4_-induced liver fibrosis model. To this end, one week after intrahepatic administration of either Lv.TβRII-SE/Fc or control Lv.TβRII-Fc, liver fibrosis was induced by chronic administration of the hepatotoxic agent CCl_4_ for 8 weeks (Fig. 5a). As controls, we used rats injected with either, CCl_4_ only, or with oil (Vehicle) also for 8 weeks. Macroscopic liver examination showed, in Vehicle rats, a reddish color, an smooth lustrous surface, and a regular shape (Fig. 5b). Conversely, livers of CCl_4_-treated animals showed irregular outlines, opaque color, shrinkage, and unsmooth surfaces. Livers of the Lv.TβRII-SE/Fc + CCl_4_ group were redder, more regular in shape, and with smoother surfaces, than CCl_4_group, and similar to the Lv.TβRII-Fc + CCl_4_ group. Also, body weight (BW) of rats was monitored throughout the experiment. Compared to the Vehicle group, CCl_4_ treatment decreased final BW gain (Fig. 5c). On the other side, BW gain was evident after 4 weeks of CCl_4_ administration, in both the Lv.TβRII-SE/Fc + CCl_4_ group and the Lv.TβRII-Fc + CCl_4_ group. In addition, CCl_4_ administration increased liver to body weight ratio (LW/BW) compared to the Vehicle group. Contrarily, Lv.TβRII-SE/Fc injected animals showed LW/BW comparable to that found in both the Vehicle and the Lv.TβRII-Fc + CCl_4_ groups (Fig. 5d).

**Figure 5.**
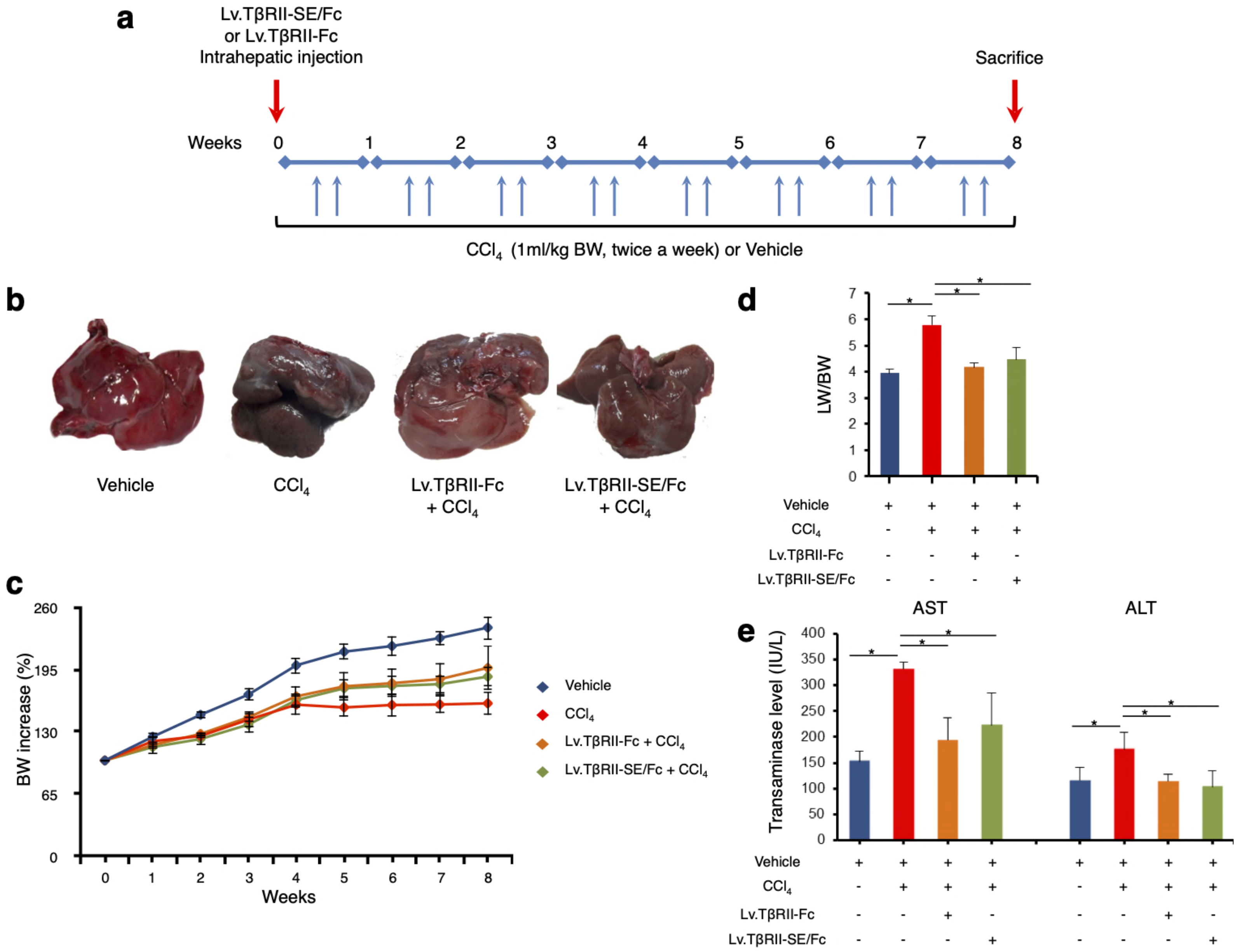
TβRII-SE/Fc overexpression prevents CCl_4_-induced liver damage development. **a** Experimental design including lentiviral vector and CCl_4_ administrations. Animals were euthanized by CO_2_ inhalation after 72 hours of the last CCl_4_ injection. **b** Representative images of liver gross appearance corresponding to rats treated with vehicle, CCl_4_, Lv.TβRII-Fc + CCl_4,_ and Lv.TβRII-SE/Fc + CCl_4_. **c** Percentage (%) of both body weight gain (left panel), and **d** liver to body weight ratio (LW/BW) (right panel) of animals in the different experimental groups. *p<0.05, Vehicle vs CCl_4_, or CCl_4_ vs Lv.TβRII-SE/Fc + CCl_4_. **e** Serum activity levels of liver enzymes aspartate aminotransferase (AST) and alanine aminotransferase (ALT) in the different experimental groups. Results are expressed as IU/L. *p<0.05, Vehicle vs CCl_4_, or CCl_4_ vs Lv.TβRII-SE/Fc + CCl_4_. IU: International units.

We also evaluated the effect of TβRII-SE/Fc on CCl_4_ induced liver injury measuring the activity level of serum aspartate aminotransferase (AST), and alanine aminotransferase (ALT). As expected, CCl_4_ administration highly increased liver enzymes above Vehicle group levels. On the other hand, Lv.TβRII-SE/Fc + CCl_4_ group showed enzyme levels comparable to both the Vehicle and the Lv.TβRII-Fc + CCl_4_ groups (Fig. 5e). These results suggested that TβRII-SE/Fc, as well as Tβ-RII-Fc overexpression, prevented CCl_4_-induced liver damage. Moreover, liver section analysis further confirmed this finding. H&E staining revealed that vehicle-injected animals presented livers with a conserved liver architecture, with cords of hepatocytes radiating from central vein (Fig. 6a). Conversely, CCl_4_ administration for 8 weeks led to a disrupted liver architecture, extensive liver injury, and prominent fibrosis. These detrimental effects were clearly attenuated when animals were injected with both Lv.TβRII-SE/Fc and Lv.TβRII-Fc, prior to CCl_4_ treatment (Fig. 6a and b). Additionally, we observed, by Sirius Red (SR) staining (Fig. 5a and b), extensive deposition of collagen fibers with bridging fibrosis in the CCl_4_ group. Similar to the Lv.TβRII-Fc + CCl_4_ group, Lv.TβRII-SE/Fc + CCl_4_ group showed reduced fibrosis located around portal areas. In addition, we evaluated hepatic stellate cells (HSC) activation by alfa-Smooth Muscle Actin (α-SMA) immunostaining (Fig. 6a and b). In this way, we observed increased α-SMA-positive areas in CCl_4_ group. Instead, HSC activation was markedly reduced in the Lv.TβRII-SE/Fc + CCl_4_ group, at levels comparable to the Lv.TβRII-Fc + CCl_4_ group. These results suggested that TβRII-SE/Fc, as well as TβRII-Fc overexpression, prevented CCl_4_-induced liver fibrosis.

**Figure 6.**
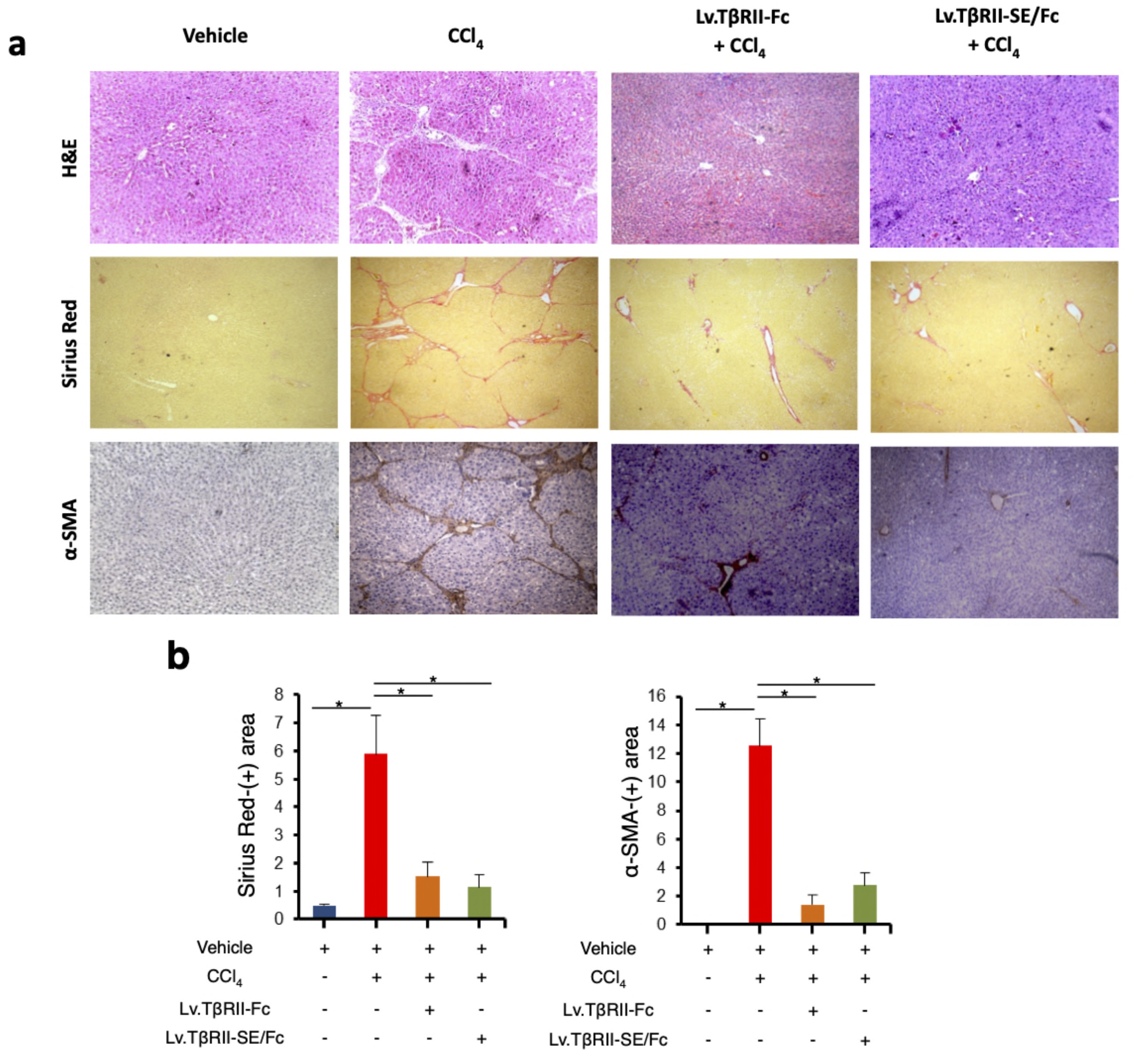
TβRII-SE/Fc overexpression prevents CCl_4_-induced liver fibrosis development and HSC activation. **a** Representative images of liver sections stained with either, H&E (top panel), Sirius Red (middle panel), and α-SMA immunohistochemistry of animals treated with vehicle, CCl_4_, Lv.TβRII-Fc + CCl_4_ or Lv.TβRII-SE/ Fc + CCl_4_. Original magnification x40. **b** Quantification of both liver Sirius Red positive areas (SR^+^) (left panel) and liver α-SMA^+^ expression (right panel) in the same experimental groups. Results are expressed as mean intensity of SR^+^ area. *p<0.05, Vehicle vs CCl_4_, or CCl_4_ vs Lv.TβRII-SE/Fc + CCl_4_. HSC, hepatic stellate cells; H&E, Hematoxylin and Eosin; α-SMA; alpha-smooth muscle actin.

Taken together, our results showed that TβRII-SE is a novel TβRII splice variant encoding a 57 AA mature peptide with distinctive attributes. This new isoform is able to bind all three TGF-β ligands with high affinity, and its Fc-tagged version modulates liver fibrosis *in vivo*.

## Discussion

In the present study, we documented for the first time the presence of a novel splice variant of the Type II TGF-β receptor in human cells. Differently to the known TβRII splice variants that encode membrane-bound receptor isoforms, the newly discovered splice variant, TβRII-SE, encodes a soluble truncated receptor with distinctive structural and binding attributes.

The TβRII–TGF-β ligand interface has been defined by X-ray crystallography. There are 11 residues in TβRII-EC involved in the binding to TGF-β3, which are well conserved in human, rats, and chicken.^26^ Although 3 out of 11 residues (Leu27, Phe30 and Asp32) remain in TβRII-SE, they do not seem to participate in the protein-protein interface with TGF-β cognate ligands. However, as revealed by our 3D binding predictions, 5 residues located in the 13-AA stretch of TβRII-SE generate a novel binding interface, allowing it to compete with TβRII for the binding with TβRI and TGF-β ligands, in the trimeric complex. This prediction was confirmed by SPR analysis, where TβRII-SE showed comparable sub-nanomolar affinity for all three TGF-β ligand complexes.

The crystal structure of the TβRII-EC/TGF-β3 complex, also revealed that ten residues contacting TβRII are identical in TGF-β1 and β3.^26^ This is consistent with their high-affinity similarity toward binding TβRII. In the low-affinity ligand TGF-β2, three interfacial positions are conservatively substituted relative to TGF-β1 and β3 (Arg25>Lys, Val92>Ile, Arg94>Lys). Thus, despite the conserved nature of the substitutions, these three residues appear responsible for the diminished affinity of TGF-β2 for TβRII.^26^ For TβRII-SE, 3D predictions indicated that five residues located in the novel 13-AA stretch were responsible for TGF-β ligand binding. In the TGF-β ligand interface, these residues contact four amino acids, instead of ten residues in the interface formed by TβRII. One of these amino acids is Arg25 in TGF-β1 and β3 interface, or Arg25>Lys substituted in TGF-β2. Our SPR results showed that this substitution was not crucial to diminish TβRII-SE binding affinity for TGF-β2, obtaining values in the picomolar range as with TGF-β1 and TGF-β3.

Cytokine receptors are conventionally produced by cells as transmembrane proteins. At the same time, most (if not all) of these receptors are also present in soluble forms.^27^ Soluble cytokine receptors, are the extracellular portions of membrane-anchored receptors. Generally, they retain the capacity to bind their ligands with similar affinity as their membrane-anchored counterparts, fulfilling different functions. Some act agonistically, others are antagonists and some are even able to accomplish both functions, depending on the biological context and stoichiometry of membrane-bound and unbound receptor.^27^ Soluble cytokine receptors are mainly generated by two mechanisms: ectodomain shedding, and alternative splicing.^27^ Ectodomain shedding is a process in which the membrane-bound receptor is cleaved in close proximity to or directly within the transmembrane domain by a protease, releasing a soluble receptor into the extracellular space. Ectodomain shedding also controls the functional availability of the cell surface TGF-β receptors and, consequently, TGF-β responsiveness. TβRI receptor, but not the TβRII, is cleaved by the transmembrane metalloprotease TACE, also known as ADAM17, in response to ERK or p38 MAPK signaling.^1^ Also, betaglycan can be cleaved to release its ectodomain that then functions sequestering TGF-β. Beta-glycan extracellular domain binds all three TGF-β ligands,^1^ with highest affinity for TGF-β2.^28^ It has been hypothesized that this high affinity may be relevant for TGF-β2, which binds TβRII with lower affinity than TGF-β1 and TGF-β3.^1^ On the other hand, soluble receptors can be generated by alternative splicing of mRNA transcripts that usually encode membrane-associated receptors.^27^ This occurs when the usual exon-intron recognition sequence is not used by the spliceosome machinery, and exons are spliced out instead. As a result, generation of frameshift in the codon sequence produces proteins that contain signal peptides and lack transmembrane domains, and are therefore secreted, rather than membrane-associated proteins.^27^ This is the case for TβRII-SE splice variant, where 149 nucleotides corresponding to parts of exon II and exon III are spliced out in the mature mRNA. The lack of this sequence generates a frameshift in the codon sequence resulting in a truncated soluble receptor with an endoplasmic reticulum signal peptide and a distinctive stretch of 13 amino acid residues in the C-terminus. Additionally, we showed by SPR that TβRII-SE binds all three TGF-βs, with high affinity (picomolar range). While betaglycan also binds all three TGF-βs, SPR analysis indicated that its soluble form has lower affinity for the three TGF-β isoforms (nanomolar range),^29^ than TβRII-SE. Although both receptors seem to have overlapping binding targets, the mechanistic role of TβRII-SE remains to be elucidated.

Fc-fusion proteins have been intensely investigated for their effectiveness to treat a range of pathologies, with several Fc-fusion drugs already in the pharmaceutical market.^30^ Fc-based fusion proteins are composed of an immunoglobin Fc domain that is directly linked to another peptide. The presence of the Fc domain markedly increases the plasma half-life of the fusion protein which prolongs therapeutic activity, due to its interaction with the neonatal Fc-receptor (FcRn),^31^ as well as to slower renal clearance.^32^ Biophysically, the Fc domain folds independently and can improve the solubility and stability of the partner molecule both *in vivo* and *in vitro*. Additionally, the Fc region facilitates its purification by using protein-G/A affinity chromatography during manufacture, in a cost-effective manner.^33^ Protein-G/A purified TβRII-SE/Fc allowed us to initially determine the apparent molecular mass of the monomer as a broad band of around 37 kDa. This observation is in agreement with the sum of the predicted molecular weight of TβRII-SE (6.5 kDa), plus the apparent molecular mass of IgG1-Fc domain (31-32 kDa) in SDS-PAGE under reducing conditions, as a result of glycosylation.^34^

Tissue fibrosis is a leading cause of morbidity and mortality worldwide affecting many organs including lung, heart, kidney, and liver.^35^ TGF-β is considered a master regulator of ECM accumulation and, consequently, a key driver of fibrosis.^36^ In the liver, TGF-β is responsible for transdifferentiation of quiescent HSC to a MFB phenotype. MFBs are characterized by the expression of α-SMA, and they are the main producers of fibrogenesis mediators and ECM proteins.^23^ Given the prominent role of TGF-β in hepatic fibrogenesis, we checked TβRII-SE functionality in CCl_4_-induced liver fibrosis rats. In this model, we observed that hepatic overexpression of TβRII-SE/Fc diminished liver damage, and attenuated liver fibrosis development to levels shown by TβRII-Fc.

Chimeric soluble TβRII (TβRII-Fc), also named soluble TβRII (sTβRII), or Fc:TβRII, has been constructed by fusion of genes encoding the N-terminal (extracellular) fragment of TβRII and the Fc-domain of human IgG. This chimeric protein has demonstrated efficacy for selective blocking of TGF-β family ligands in pathological conditions,^37,38^ including liver fibrosis.^39^ However, its therapeutic potential is isoform-selective, as TGF-β1 and TGF-β3 are known to bind TβRII receptor dimers with high affinity, but not TGF-β2,^9^ as we confirmed here by SPR analysis. TGF-β family proteins are secreted and function as disulfide-linked homodimers or heterodimers,^1^ therefore the stoichiometry of TGF-β dimer to TβRII dimer is known to be 1:1. Our SPR analyses confirmed this observation for TβRII-Fc and indicated the same stoichiometry for TβRII-SE/Fc.

Here we also show that TβRII-SE/Fc is as effective at binding all three TGF-β isoforms as 1D11 mAb. This is a mouse pan-TGF-β neutralizing antibody,^40^ that is the parent antibody of the humanized and optimized version GC-1008.^41^ The crystal structure of GC-1008 in complex with TGF-β3 showed that all CDR loops of the heavy chain are involved in the binding of TGF-β as well as some residues of the FWH3. The Fab recognizes parts of both TGF-β3 molecules in the homodimer, not only residues from a single monomer.^42^ The total binding area of TGF-β to the antibody consist of 2 identical binding interfaces on the surface of the TGF-β3 homodimer. Our binding data confirmed that 1D11 binds a dimer per Fab as reflected in the calculated stoichiometry of TGF-β homodimer per antibody of 2:1.

In this article we demonstrate that TβRII-SE/Fc and TβRII-Fc share similarities, such as binding affinity to TGF-β isoforms, *in vitro* blockade of TGF-β1 signaling, and *in vivo* prevention/inhibition of liver fibrosis. Thus, we hypothesized that, like TβRII-Fc, TβRII-SE/Fc would be able to act as a ligand trap, despite further studies must be conducted to clarify its mechanism of action.

Although TGF-β1 is the best characterized pro-fibrotic factor within the family, TGF-β2 also displays potent fibrotic activity. TGF-β2 is increased in late stages of liver fibrosis associated to hepatitis C virus (HCV) infection in patients;^43^ is accumulated in the bile ducts in human fibrotic liver disease;^44^ and has been implicated in the fibrotic response associated with glaucoma.^45^ Information regarding TGF-β3 is scarce in the literature. *In vitro*, TGF-β3 has been reported to exert profibrotic effects in fibrosis-related cell types,^46,47^ such as fibroblasts. Thus, the fact that TβRII-SE binds all three TGF-β ligands with high affinity could offer an alternative strategy to treat fibrosis related diseases. Also, increased expression of all three biologically active TGF-β isoforms has been reported in various cancers including glioblastoma, breast cancer, and colorectal cancer. In particular, high TGF-β2 expression is associated with poor prognosis of advanced lung cancer, gliomas, and skin squamous carcinoma.^48^ Therefore, TβRII-SE/Fc may be a promising tool for the prevention and treatment of pathological conditions caused by TGF-β upregulation, especially TGF-β2.

## Materials and methods

### Cell Culture

Human adipose derived mesenchymal stromal cells (hASC), purified as described,^49^ and human cell lines A549, Jurkat, 293T and HCT116 were cultured in DMEM supplemented with 10% FBS and 1% penicillin/streptomycin, in a humidified 5% CO_2_ incubator at 37°C.

### Purification of leukocyte subsets

Granulocytes, lymphocytes and monocytes were isolated from heparinized human peripheral blood by Ficoll-PaqueTM PLUS (GE Healthcare Bio-Sciences AB, Uppsala, Sweden) gradient centrifugation. After centrifugation, two fractions were obtained: one containing granulocytes/erythrocytes and another with peripheral blood mononuclear cells (PBMCs). To obtain granulocytes, erythrocytes were lysed with KCl 0,6 M. PBMCs were labelled with anti CD3^+^, CD14^+^, and CD19^+^ monoclonal antibodies conjugated with magnetic microbeads (Miltenyi Biotech, Bergisch Gladbach, Germany) and separated using MS columns in a MiniMACS magnet (Miltenyi Biotech, Bergisch Gladbach, German). Viable cells were determined by Trypan blue dye exclusion and counted in a hemocytometer. The purity of CD19^+^, CD3^+^ and CD14^+^ cells was determined by flow cytometric analysis using a FACSCalibur flow cytometer (BD Biosciences, San Jose, CA). Cell sub-populations were homogenized in RNA Lysis Buffer (SV Total RNA Isolation System, Promega Corporation, Madison, WI) and stored at -80°C until RNA purification.

### End point RT-PCR

Total RNA from different primary cultures and cell lines was isolated using the SV Total RNA Isolation System, and cDNA was generated using 1 mg RNA, M-MLV Reverse Transcriptase, and oligo dT_(15)_ primers, according to the indications stated by the manufacturer (Promega Corporation, Madison, WI). To simultaneously detect the different splice variants of the TβRII receptor, PCR amplification was performed in the presence of Expand High Fidelity polymerase (Roche Diagnostics GmbH, Mannheim, Germany), 0,2 mM dNTPS, and 0,5 μM of each primer (forward: 5’ACCGGTATGGGTCGGGGGCTGCTC3’ and reverse: 5’gtcgactcagtag CAGTAGAAGATG3’ for 35 cycles using the following PCR conditions: 1 min. at 95°C, 1 min. at 55°C, and 1 min. at 95°C.

### DNA sequencing

PCR fragments were cloned into the pGEM-T Easy plasmid (Promega Corporation, Madison WI) under the conditions established by the manufacturers and *E. coli* transformation. TβRII PCR fragments were sequenced by using M13 forward and reverse primers using a capillary automatic sequencer model ABI3130XL (Applied Biosystems, USA) at the Genomic Unit of the Biotechnology Institute, INTA, Consorcio Argentino de Tecnología Genómica (CATG) (PPL Genómica, MINCyT).

### TβRII-SE codon optimization and human IgG1 Fc chimeric fusion protein

TβRII-SE was codon optimized (co) together with the deletion of the stop codon. This product also included a Kozak sequence, and an *Age*I site and *EcoR*V at the 5’ and 3’ end, respectively (Epoch Biolabs Inc., Missouri City, TX). The human IgG1 Fc coding sequence was obtained by RT-PCR from peripheral blood leucocyte mRNA using specific oligonucleotides as primers (forward: 5’-AGATCT-GACAAAACTCACACATGC-3’ and reverse:5’-GATATCTTTACCCGGAGACAGG-3’), containing a *Bgl*II recognition site (forward primer) and an *Eco*RV site (reverse primer). The coTβRII-SE sequence (258 bp) was fused *in frame* with the coding sequence of the IgG1 Fc domain (693 bp) to generate the coTβRII-SE/Fc fusion cDNA of 951 bp.

### Lentiviral vector production

The cDNA encoding TβRII-SE with IgG1 Fc, was cloned into the pRRLsin18.cPPT.WPRE lentiviral vector, generating the vector pRRLsin18.cPPT.CMV-coTβRII-SE/Fc.ires.eGFP.WPRE (Lv.TβRII-SE/Fc). As control, we cloned the coding sequence of TβRII extracellular domain (TβRII-EC) fused in frame also with IgG1 Fc to generate the vector pRRLsin18.cPPT.CMV-Tβ-RII-Fc.ires.eGFP.WPRE (Lv.TβRII-Fc). Vesicular Stomatitis Virus G protein-pseudotyped lentiviruses (VSV-G) were generated by transient transfection of the transfer vectors together with the envelope plasmid (pCMV-VSVG), the packaging plasmid (pMDLg/pRRE) and Rev plasmid (pRSV-REV), into the 293T cell line, as previously described.^50^ Cell supernatants were harvested once every 12 hours, for 48 hours, and frozen in aliquots. Viral titers were determined by transducing A549 cells, yielding 10^7^ TU (transducing units) per milliliter.

### Smad2/3 activation and immunoblotting

Human colorectal cancer-derived cell line HCT116 cells were transduced with Lv.TβRII-SE/Fc at MOI 200, in the presence of 8 μg/ml polybrene (Millipore Sigma, Burlington, MA). HCT116 cells (1 x 10^6^) overexpressing TβRII-SE/Fc or control were seeded in 60 mm cell culture dishes and starved for 24 hours in DMEM. After that, cells were incubated in DMEM ± 5 ng/mL TGF-β1 for 1 hour. Cells were lysed in RIPA buffer (Millipore Sigma, Burlington, MA) supplemented with 1 mM PMSF and quantified by Bradford Assay. Proteins were separated by electrophoresis on 10% SDS-PAGE gels and electrotransferred onto Immobilon-polyvinylidene difluoride membranes (Millipore Sigma, Burlington, MA). After blocking with 5% non-fat milk, membranes were probed with antibodies for P-Smad2/3 (sc-11769) and Smad2/3 (sc133098) (Santa Cruz Biotechnology, Inc., Dallas, TX) at 4°C overnight and then incubated with horseradish peroxidase (HRP)-conjugated secondary anti-mouse or rabbit antibodies (Thermo Fisher, Waltham, MA). Protein expression was detected by using enhanced chemiluminescence (ECL) system (Thermo Fisher, Waltham, MA). Densitometry was performed using ImageJ software (National Institutes of Health, Bethesda, MD).

### TβRII-SE/Fc-fusion recombinant protein production and purification

Human 293T cells were transduced either with Lv.TβRII-SE/Fc or Lv.TβRII-Fc at a MOI of 70, in the presence of 8 μg/ml polybrene (Millipore Sigma, Burlington, MA). Forty-eight hours after transduction, cells were harvested, washed in phosphate-buffered saline (PBS) supplemented with 10% FBS, and the percentage of eGFP positive cells was determined by flow cytometry in a FACSCalibur device (BD Biosciences, San Jose, CA).

For the production of TβRII-SE/Fc and TβRII-Fc, lentivirally transduced 293T cells were cultured for 48 h in serum-free DMEM supplemented with Protease Inhibitor Cocktail (1/800) (Millipore Sigma, Burlington, MA). Subsequently, the conditioned media was clarified by centrifugation at 3500 rpm and filtrated through 0.22 μm filters. Recombinant protein was concentrated by centrifugation in Amicon® Ultra-15-30K (Millipore Sigma, Burlington, MA), purified on protein A/G columns (NAb™ Spin Kit, Thermo Scientific, Rockford, IL) following instructions of the manufacturer, and stored at -80°C.

### TβRII-SE 3D modeling and molecular dynamic simulation

We generated a preliminary 3D model of the TβRII-SE peptide structure using Robetta server.^51^ Robetta produces protein models using comparative modeling and *ab initio* methods. Domains without a detectable template are modeled using Rosetta *de novo* algorithm. In the initial stage, Robetta identified the extracellular (EC) domain of human TβRII (Protein Data Bank of the Research Collaboratory for Structural Bioinformatics (RCSB-PDB): 1PLO_A) as a confident template for TβRII-SE modeling using the method “Ginzu”, and used it as a template to produce models with a comparative modeling protocol. Robetta carried out multiple independent simulations to generate thousands of models, and finally, five of them were selected from this ensemble applying different variants of the Rosetta energy function. We selected the best model with lowest energy informed by Robetta to be used in further analyses. We selected the best model with the lowest energy informed by Robetta and proved its stability along a 1000 ns long molecular dynamics trajectory during which the model maintained its fold and Secondary Structure Elements. Thus, we used this model for further analyses (see Supplementary Materials and methods for further details).

To predict the binding of TβRII-SE to its TGF-β ligands, we used the ternary complexes of the structures PDB code: 3KFD (TβRII/TβRI/TGF-β1) and PDB code: 2PJY (TβRII/TβRI/TGF-β3) deposited in the Protein Data Bank (PDB), as templates. In addition, TβRII-SE/Fc fusion protein was modeled using the structures PDB code: 2PJY (TβRII) and PDB code: 1L6X (Fc), as templates.

### Surface plasmon resonance (SPR) assays

SPR experiments were performed using a Biacore 3000 system (Cytiva Life Sciences, Marlborough, MA). All assays were performed using CM5 sensor chips (Cytiva Life Sciences, Marlborough, MA) derivatized as described previously,^52^ with anti-hu Fc mixed with equimolar anti-mu Fc antibodies (cat # 109-005-008 and 115-005-008, respectively; Jackson ImmunoResearch, West Grove, PA) at pH 5.0, obtaining four ∼6000 RU anti-Fc surfaces including reference surface or non-specific binding control. Human Fc recombinant proteins TβRII-SE/Fc (64 kDa), TβRII-Fc (124 kDa) and murine mAb 1D11 (150 kDa) (Thermo Fisher, Waltham, MA) were each captured on individual flow cells to 100-140 RU. Captures were designed with a wait time of 60 s after the association to reach a stable signal, and the capture antibodies reference flow cell had no ligand captured. TGF-β1, TGF-β2 and TGF-β3 (PeproTech, Cranbury, NJ), were individually injected over the four flow cells over range of concentrations prepared by serial 2-fold or 3-fold dilutions, spanning from 0.4 to 30 nM, or at two, 5-fold apart, concentrations in higher throughput tests depending on the experimental design, all at a flow rate of 50 μl min^-1^. Non-specific binding of the TGF isoforms to the reference capture antibody surfaces was initially detected and minimized by increasing 20-fold the surfactant concentration in the sample and running buffers from 0.005% to 0.1% as previously recommended.^53^Multiple-cycle kinetics were programed using an association time of 90 seconds and dissociation time of 300 seconds. Capture surfaces were regenerated using three 3s, 100 mM phosphoric acid (pH 2.0). Assessment of dissociation for 900 s allowed to confirm 10% loss of complex to measure the slower dissociation rates to calculate the dissociation constants (*k*_*d*_). Running buffer samples were also injected using the same method program for background noise and capture level drift subtraction (double reference). All data were fitted to a 1:1 binding model using Biacore Evaluation Software 3.1 (Cytiva Life Sciences, Marlborough, MA) and Scrubber 2.0c (BioLogic Software, Australia) after double referencing, where n = 6 independent experiments using 3 capture chips, presenting kinetic constants ±SEM, and thermodynamic constant K_D_ ±SEM, 95% CI (confidence interval). Mass ratios of approximately 2.5:1, 5:1 or 6:1 for TβRII-SE/ Fc, TβRII-Fc and 1D11, respectively, were used to estimate stoichiometries of complexes formation.

### Animals

Male Wistar rats (150-200 g) were housed at Mar del Plata National University Laboratory Animal Unit at a mean constant temperature of 22°C with a 12 h light– dark cycle, and free access to standard pellet chow and water. All experiments were performed according to the ‘Guide for the Care and Use of Laboratory Animals’ and approved by the Institutional Animal Care and Use Committee (CICUAL) of Mar del Plata National University.

### *In vivo* liver transduction and liver fibrosis induction

Liver fibrosis was induced by intraperitoneal injection (i.p.), of 1 ml/kg body weight CCl_4_ in oil (1:1) twice a week for 8 weeks. A week before liver fibrosis induction, rats received an intrahepatic injection of either Lv.TβRII-SE/Fc or Lv.TβRII-Fc (5-10 x 10^7^ TU/ml) (N=6 each group). Lentiviral vectors were directly injected, after a small incision in the abdomen, into exposed livers of anesthesized rats. Control animals received i.p. injection of either CCl_4_ or oil (vehicle) (N=6 each group). Animals were weekly weighted, and measurements were used to calculate the percentage of body weight gain. After eight weeks, animals were euthanized, and livers were weighted to calculate liver to body weight ratio (LW/BW). Livers were fixed in 10% neutral buffered formalin for histological analysis. Serum was also collected for further biochemical analyses.

### Biochemical parameters

Serum Aspartate aminotransferase (AST) and Alanine aminotransferase (ALT) activity levels were measured with an automatic analyzer BT300 plus (Biotecnica Instruments S.p.A., Rome, Italy) according to the manufacturer.

### Histological analysis

Livers fixed in 10% neutral buffered formalin were embedded in paraffin. Liver sections (5 µm) were stained with Hematoxylin and Eosin staining (H&E) for liver architecture visualization. To assess fibrosis, liver sections were stained with Sirius Red staining (0.1%). Quantification of Sirius Red-positive (SR^+^) areas was performed in at least ten fields per histological section using the ImageJ software. Results were expressed as mean intensity of SR^+^ area per field.

### Immunohistochemichal analysis

For immunohistochemical analysis, 5-µm sections were deparaffinized and rehydrated. Endogenous peroxidase was blocked through the addition of 3% H_2_O_2_, in methanol. Antigen retrieval was performed using heat induced epitope retrieval (HIER) method with citrate buffer 0.1 M, pH 6. Tissue sections were then incubated with rabbit anti-α-smooth muscle actin (anti-α-SMA, 1:500, Cell Signaling Technology, Danvers, MA) overnight at 4°C. After two washes with PBS, slides were incubated with HiDef Detection amplifier Mouse and Rabbit reagent (Cell Marque, Rocklin, CA) for 10 min, at room temperature. Sections were further washed with PBS and incubated with HiDef Detection HRP Polymer Detector solution (Cell Marque, Rocklin, CA) for 10 min., at room temperature. Finally, sections were washed twice with PBS, incubated with DAB Chromogen kit (Cell Marque, Rocklin, CA) for 5 min. at room temperature, and counterstained with Hematoxylin. Dehydrated sections were mounted, and microphotographed on a light microscope Nikon Eclipse E200. Quantification of α-SMA-positive (α-SMA^+^) areas was performed through use of Fiji software. Results were expressed as mean intensity of α-SMA^+^ area per field.

### Statistical analysis

Statistical analysis was performed through GraphPad Prism Version 7.0 (GraphPad Software, San Diego, CA). Data are shown as mean ± SD. Statistical differences among groups were performed using two-way ANOVA and the multiple comparison post-hoc test by Fisher. For all analyses, a P-value < 0.05 was considered statistically significant.

## Data availability

All the data that support the findings of this study are available within the article or from the corresponding author upon reasonable request.

## Acknowledgments

We wish to acknowledge Professor Isabel Fabregat at Bellvitge Biomedical Research Institute (IDIBELL), Barcelona, Spain, and Professor Irwin Chaiken at the Biochemistry and Molecular Biology Department from Drexel U Medical College, USA, and Director of the S200 Biosensor Shared Resource, for their critical reading of the manuscript and useful suggestions. This work was supported, in part, by the National Scientific and Technical Research Council of Argentina (CONICET) research grants PIP 2013 GI, 11220120100202CO and PIP 2017 GI, 11220170100573CO, by additional financial support from Fundación Articular, (Quilmes, Buenos Aires, Argentina), Fundación Florencio Fiorini (Academia Nacional de Medicina, Buenos Aires, Argentina), and the startup RAD BIO S.A.S. (Sunchales, Santa Fe, Argentina).

## Conflict of Interest

G. P. Barletta, A.M. Monzón, S.M. Echarte, S. Campisano and G. Canziani declare no conflict of interest. A. Carrea, T.M. Rodríguez, A.N. Chisari and R.A. Dewey are co-inventors of the patent family “Isoform of the TGF-beta receptor II”, US10233227B2 (granted in USA), EP3082846B1 (granted by the European Patent Office), ES2749615T3 (granted in Spain) and AR098827A1 (pending in Argentina). T.M Rodríguez, A.N Chisari, A. La Colla, M.S. Bertolio, A. Romo and R.A. Dewey are co-inventors of the patent application “TGF-β receptor II isoform, fusion peptide, methods of treatment and methods *in vitro*”, US20190112352A1 (pending in USA). Patents are own by CONICET and Fundación Articular, and were licenced to RAD BIO S.A.S by Intelectual property licence agreement 2019-890-APN-DIR#CONICET. A. Romo has shareholder equity of RAD BIO S.A.S., and R.A. Dewey is co-founder, and shareholder of RAD BIO S.A.S.

## Notes

### Summary of Updates

Figure 4 revised Figure 4 caption updated Section on Results, Subsection "TβRII-SE/Fc binds with high affinity to all three TGF-β isofoms" updated to clarify SPR results

