## Supplemental material for "A novel splice variant of human TGF-β type II receptor encodes a soluble protein and its Fc-tagged version prevents liver fibrosis *in vivo*"

**This PDF file includes:**

Materials and methods
Figure. S1
Table. S1
Table. S2
Supplementary Text

### Materials and methods

#### Molecular dynamic simulation

In order to evaluate our TβRII-SE 3D model under physiological conditions, we performed a molecular dynamic (MD) simulation on the best model using the AMBER 16 software package.[1,2] Ions were added for charge neutralization. Each system was solvated with explicit TIP3P[3] water molecules in a truncated octahedral periodic box large enough to contain the protein and 10 Å of solvent on all sides. The all-hydrogen topology was obtained with the Amber ff14SB[4,5] force field. A 2 fs time step was used and the SHAKE algorithm was applied to all bonds involving hydrogen. Periodic boundary conditions and particle-mesh Ewald (PME) sums were used and a cut-off of 10 Å was applied to non-bonded interactions.

Minimization was performed by 100-step of steepest-descent and 400-step conjugate gradient minimizations applying constraints to the protein atoms. This was followed by a 400-step unconstrained conjugate gradient minimization.

The system was then heated for 150 ps until it reached the final temperature of 300 K. During heating, a harmonic restraint of 50.0 kcal/(mol·Å2) was applied to the protein atoms.

The system was equilibrated at constant pressure using 29 steps of 100 ps and reducing the restraint on each step. After the last step with restraints, all restraints were lifted and a final equilibration run was performed until reaching 5 ns of total equilibration time at a constant temperature of 300 K using Andersen barostat and Langevin thermostat with a γ collision frequency of 2 ps−1. Finally, a 1000 ns MD simulation is performed, collecting equilibrated configurations at 10 ps intervals.

**Figure. S1.**

TβRII-SE ACCGGT------ATGGGTCGGGGGCTGCTCAGGGGCCTGTGGCCGCTGCACATCGTCCTGTGGACGCGTATCGCCAGCAC
coTβRII-SE ACCGGTGCCACCATGGGAAGAGGTCTCCTCAGAGGACTCTGGCCACTGCACATCGTCCTGTGGACCAGAATCGCATCTAC

TβRII-SE GATCCCACCGCACGTTCAGAAGTCGGTTAATAACGACATGATAGTCACTGACAACAACGGTGCAGTCAAGTTTCCACAAC
coTβRII-SE CATCCCTCCTCATGTGCAGAAATCTGTCAACAATGACATGATCGTCACAGACAACAACGGTGCTGTGAAGTTTCCTCAGC

TβRII-SE TGTGTAAATTTTGTGATGTGAGATTTTCCACCTGTGACAACCAGAAATCCTGCTTCTCCAAAGTGCATTATGAAGGAAAA
coTβRII-SE TGTGTAAGTTCTGCGACGTCAGGTTCAGTACCTGCGACAATCAGAAATCTTGTTTCAGCAAGGTGCACTACGAAGGGAAG

TβRII-SE AAAAAAGCCTGG*TGA*
coTβRII-SE AAGAAAGCATGGAGATCT

**TβRII-SE codon optimization.** cDNA alignment showing changes made to TβRII-SE (yellow shading). To get coTβRII-SE, a Kozak consensus sequence (red nucleotides) was included. Additionally, some nucleotides have been substituted to make translation more efficient in human cells. To allow fusion in frame of cDNA with the human IgG-Fc domain cDNA, the stop codon of TβRII-SE was removed (italics) and replaced by a *Bgl*II recognition sequence in the new construct.

| | | |
|---|---|---|
| **pI/Mw** | 9.64/9161.72 Da | |
| **Signal peptide clivage site** | Between T23-I24 | SignalP 4.1 Server [6]<br>http://www.cbs.dtu.dk/services/SignalP/ |
| **pI/Mw without signal peptide** | 9.05/6532.51 Da | |
| **Glycation** | K46, 52, 78 | NetGlycate 1.0 Server [7]<br>http://www.cbs.dtu.dk/services/NetGlycate/ |
| **Kinase-specific Phosphorylation** | S22, 31, 59, 66, 69<br>T18, 23, 39, 60,<br>Y73 | GPS web server 5.0 [8]<br>http://gps.biocuckoo.org/online.php |
| **Sumoylation** | K76, 77, 78<br>(Non consensus) | GPS-SUMO 2.0 Online Service [9]<br>http://sumosp.biocuckoo.org/online.php |
| **C-mannosylation** | No sites | NetCGlyc 1.0 Server [10]<br>http://www.cbs.dtu.dk/services/NetCGlyc/ |
| **N-Glycosylation** | No sites | NetNGlyc 1.0 Server<br>http://www.cbs.dtu.dk/services/NetNGlyc/ |
| **GalNAc O-glycosylation** | No sites | NetOGlyc 4.0 Server [11]<br>http://www.cbs.dtu.dk/services/NetOGlyc/ |
| **N-terminal Myristoylation** | No sites | Myristoylator [12]<br>https://web.expasy.org/cgi-bin/myristoylator/myristoylator.pl |
| **Palmitoylation** | No sites | CSS-Palm 2.0 [13]<br>http://csspalm.biocuckoo.org/online.php |

**Table. S1.**

**TβRII-SE predicted characteristics including post-translational modifications.**

**Table. S2.**

| Ligands | Surface Density (RU/Kda) | Theoretical $R_{max}$ | Experimental $R_{max}$ | Stoichiometry experimental/ predicted |
|---|---|---|---|---|
| **TβRII-SE/Fc** | 2.0 | 60.0 | 65.7 ± 12.9 | 1.1/1 |
| | 2.5 | 62.5 | 59.6 ± 9.3 | 0.95/1 |
| | 3.0 | 90.0 | 59.2 ± 7.3 | 0.95/1 |
| **1D11** | 0.8 | 80.0 | 126 ± 4.5 | 2.5/2 |
| | 1.0 | 100.0 | 120 ± 6.0 | 2.4/2 |
| | 2.4 | 120.0 | 60.8 ± 9.9 | 1.0/2 |
| **TβRII-Fc** | 0.9 | 22.5 | 44.5± 1.9 | 2.0/1 |
| | 1.0 | 25.0 | 48.2 ± 1 | 1.9/1 |
| | 1.2 | 30.0 | 57.2 ± 2.3 | 1.9/1 |

**Stoichiometries of binding of TGF-β dimers to their experimental ligands.**

**Supplementary Text**
